# Dynamic tracking of objects in the macaque dorsomedial frontal cortex

**DOI:** 10.1101/2022.06.24.497529

**Authors:** Rishi Rajalingham, Hansem Sohn, Mehrdad Jazayeri

## Abstract

Cognitive neuroscience posits that humans perform physical inferences using mental simulations. Here, we test this hypothesis by analyzing neural activity in the monkeys’ frontal cortex in a ball interception task. We find a low-dimensional neural embedding of the ball position that tracks the ball both when it is visible and invisible. This embedding may serve as a neural substrate for mental simulation.

## Introduction

With just a few glances, humans can make rich inferences about objects and predict their future states. For example, we can predict how balls would move around a billiard table from random initial configurations ^1^, or the probability that a perturbation would tear down a tower of blocks ^2^. A dominant cognitive theory posits that humans make these predictions by forming mental models of the physical world and using those models to perform simulations ^3–5^. In support of this theory, it has been shown that human predictions are consistent with simulations of abstract models instantiated by high-level computer programs ^2,6^. However, evidence for the defining feature of mental simulation – that the brain reproduces the neural states associated with physical experiences – is lacking ^7,8^. Tackling this question in the brain and at the level of neural populations is critical for two reasons. First, artificial neural networks can perform mental computations with dynamics that differ widely from those generated during physical experiences ^9–12^. Second, neuroimaging studies in humans that have attributed elevated activity in various brain areas to mental simulations do not have the resolution needed to verify that the underlying brain dynamics are indeed consistent with such simulations ^13–16^. In light of these considerations, it is important to revisit the mental simulation hypothesis in two ways. First, we need to show that neural network models whose dynamics are consistent with simulations are superior to other models with arbitrary dynamics in terms of capturing behavior. Second, we need to record directly from populations of neurons in candidate brain areas to verify whether and when brain dynamics during mental computations are consistent with the simulation hypothesis.

We recently addressed the first of these two desiderata in a behavioral/modeling paper by comparing the behavior of humans and monkeys (i.e., “primates”) to that of recurrent neural network (RNN) models in a ball interception task in which the ball trajectory was partially occluded (“Mental-Pong”). Consistent with the mental simulation hypothesis, we found that RNNs trained to track the instantaneous position of the occluded ball (i.e., “simulate”) generated behavioral patterns that were more similar to primates than models that were not constrained as such ^10^. Here, we use monkey electrophysiology to address the second and arguably more critical prediction that the underlying neural signals in candidate brain areas are consistent with simulation of the latent position of the occluded ball.

## Results

We devised a virtual task in which subjects have to use a joystick to control the vertical position of a paddle with the goal of intercepting a ball moving across a two-dimensional frame with reflecting horizontal “walls” (Figure 1A). The ball’s initial position and velocity (speed and heading) are randomly sampled on every trial with the constraint that the ball reaches the end point within a given time window either along a straight line or after bouncing off of the top or bottom wall (see Methods). Importantly, the frame contains a large occluder covering all trajectories before the interception point. Therefore, the ball is visible only during the first portion of the trial and invisible afterwards. We refer to this task as Mental-Pong because of its similarity to the computer game Pong, and because of the presence of the occluder that necessitates mental (as opposed to visually-driven) computations.

**Figure 1.**
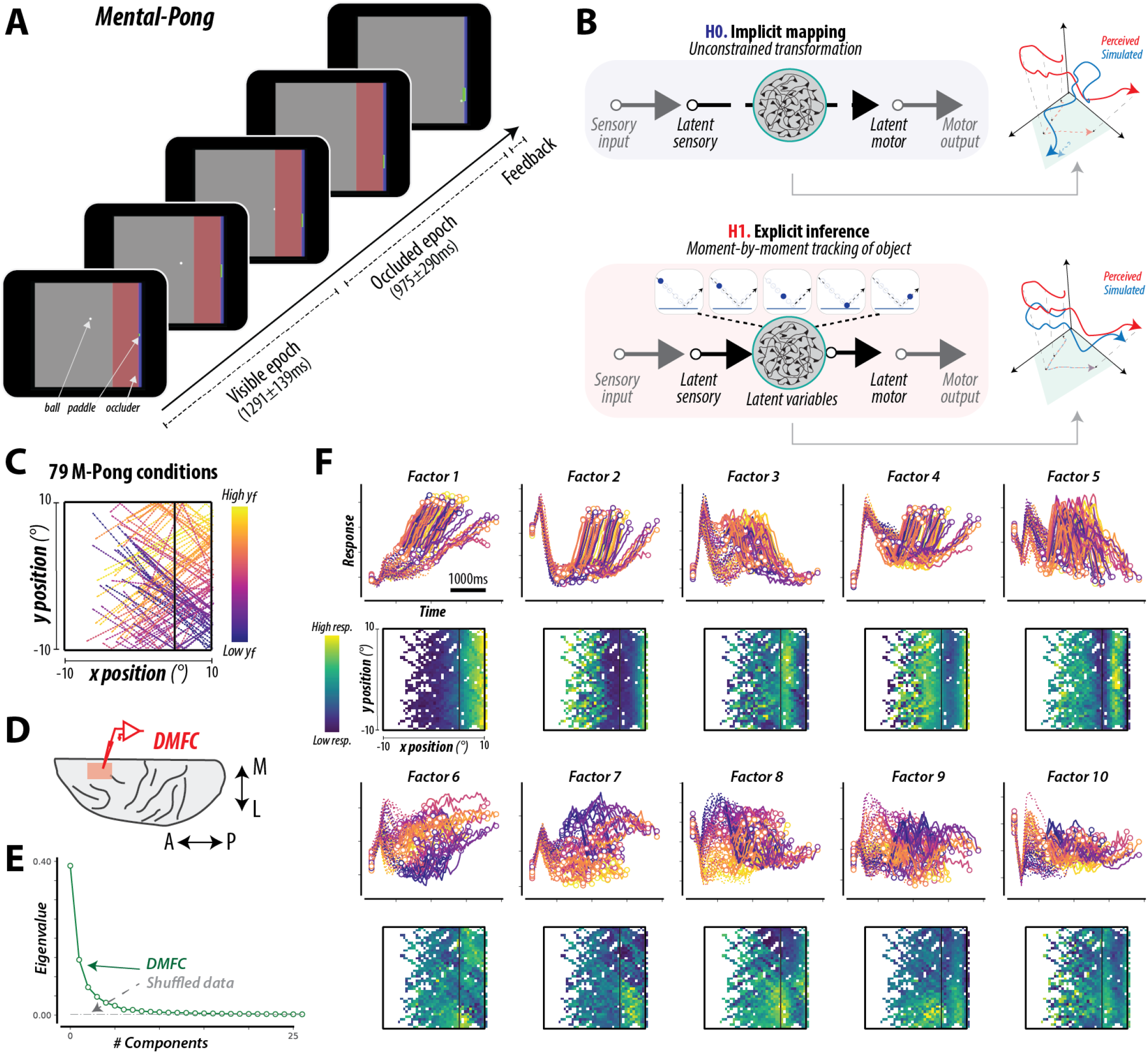
**(A) Behavioral task.** The time-course of a trial of Mental-Pong is shown. The objective of the task is to intercept a ball moving across a two-dimensional frame with reflecting walls, by manipulating a joystick to control the vertical position of a paddle. The ball’s initial position and velocity were randomly sampled on every trial. Moreover, the frame contained a large rectangular occluder before the interception point such that the ball’s trajectory was visible only during the first portion of the trial. Monkeys initiated each trial by fixating on a central fixation dot, but were subsequently free to make any eye movements. On every trial, monkeys could move the paddle as soon as the ball began to move, and could drive it freely and at a constant speed in up or down directions from its initial position at the middle of the screen with the goal of intercepting the ball when it reached the paddle. We refer to this task as Mental-Pong because of its similarity to the computer game Pong, and because of the presence of the occluder that necessitates mental (as opposed to visual) computations. **(B) Computational hypotheses**. The top left panel illustrates the implicit mapping hypothesis, whereby latent sensory representations are transformed via nonlinear functions (denoted via an RNN), without explicitly tracking latent environmental states. The bottom left panel illustrates an explicit inference strategy, whereby specific neural representations dynamically track latent environmental states to generate appropriate motor outputs. The right panels schematically illustrate the internal activity representations for each of these computational hypotheses. Explicit inference implies that the brain reproduces the neural states associated with physical experiences, such that activity related to “perceived” and “simulated” experiences are similar along specific dimensions. **(C)** The ball trajectories of the 79 unique conditions used for this study; colors correspond to the per-condition endpoint y-position of the ball (yf). **(D)** Neurophysiology targeted the dorsomedial frontal cortex (DMFC). **(E)** Neural dynamics in DMFC were relatively low-dimensional, as quantified by the eigenspectrum, estimated using PCA. The dashed gray curve corresponds to the eigenspectrum estimated after shuffling the data matrix. **(F)** Neural dynamics in DMFC, where the activity of 1889 neurons is summarized via ten factors. For each factor, the top panels show the time-locked response for each of the 79 Mental-Pong conditions (dotted lines for visible epoch, solid lines for occluded epoch, circular markers for epoch boundaries); color mapping identical to the panel (C). The bottom panels show the same factor activity as a heat map over the visuospatial dimensions. The boundary of the Mental-Pong frame is denoted in black.

Our key question of interest is whether the brain’s activity during the time when the ball is invisible is consistent with simulating the latent ball position. If the brain were to perform such a simulation, we would expect the underlying neural responses during occlusion to have dynamics similar to when the ball is visible. We refer to this strategy as *explicit inference* (Figure 1B, *H1*). However, solving Mental-Pong does not necessitate mental simulation. As previous work has demonstrated ^10^, a recurrent neural network can compute the endpoint through dynamics that do not correspond to the moment-by-moment latent position of the occluded ball. We refer to this alternative strategy as *implicit mapping* (Figure 1B, *H0*).

The key difference between these hypotheses is that, under explicit – not implicit – inference, the neural signals should represent the moment-by-moment position of the ball, especially while the ball is moving behind the occluder. To test these predictions, we focused on the dorsomedial frontal cortex (DMFC) that has been implicated in mental simulation ^13^ and where neural signals have complex heterogeneous dynamics ^17,18^. We recorded from a large population of individual neurons (1889 neurons; 1552 in monkey P, 337 in monkey M) while monkeys performed 79 conditions of Mental-Pong (Figure 1C,D, S1A). Consistent with previous work ^17,18^, neurons had heterogeneous and complex response profiles with reliable spatiotemporal properties (Figure S1B). However, across neurons, DMFC population dynamics could be explained in terms of a small number of latent factors (Figure 1E), each exhibiting rich and reliable spatiotemporal dynamics (Figure 1F).

Next, we asked if DMFC has an explicit representation of the instantaneous ball position. To address this question, we constructed linear decoders of the instantaneous ball position [x(t), y(t)] from the latent factors (Figure 2A). We reasoned that if DMFC carries such an explicit representation, then a single decoder should be able to extract [x(t), y(t)] for the entire duration of the trials and that the decoder should generalize across conditions. Results were consistent with the presence of an explicit representation of the ball’s position: a single decoder accurately predicted the instantaneous ball position (Figure 2B), robustly generalized across conditions with and without bounces (Figure S2A), and was temporally stable across visible and occluded epochs (Fig 2C, 2D). These results were consistent across both monkeys (Figure S3).

**Figure 2.**
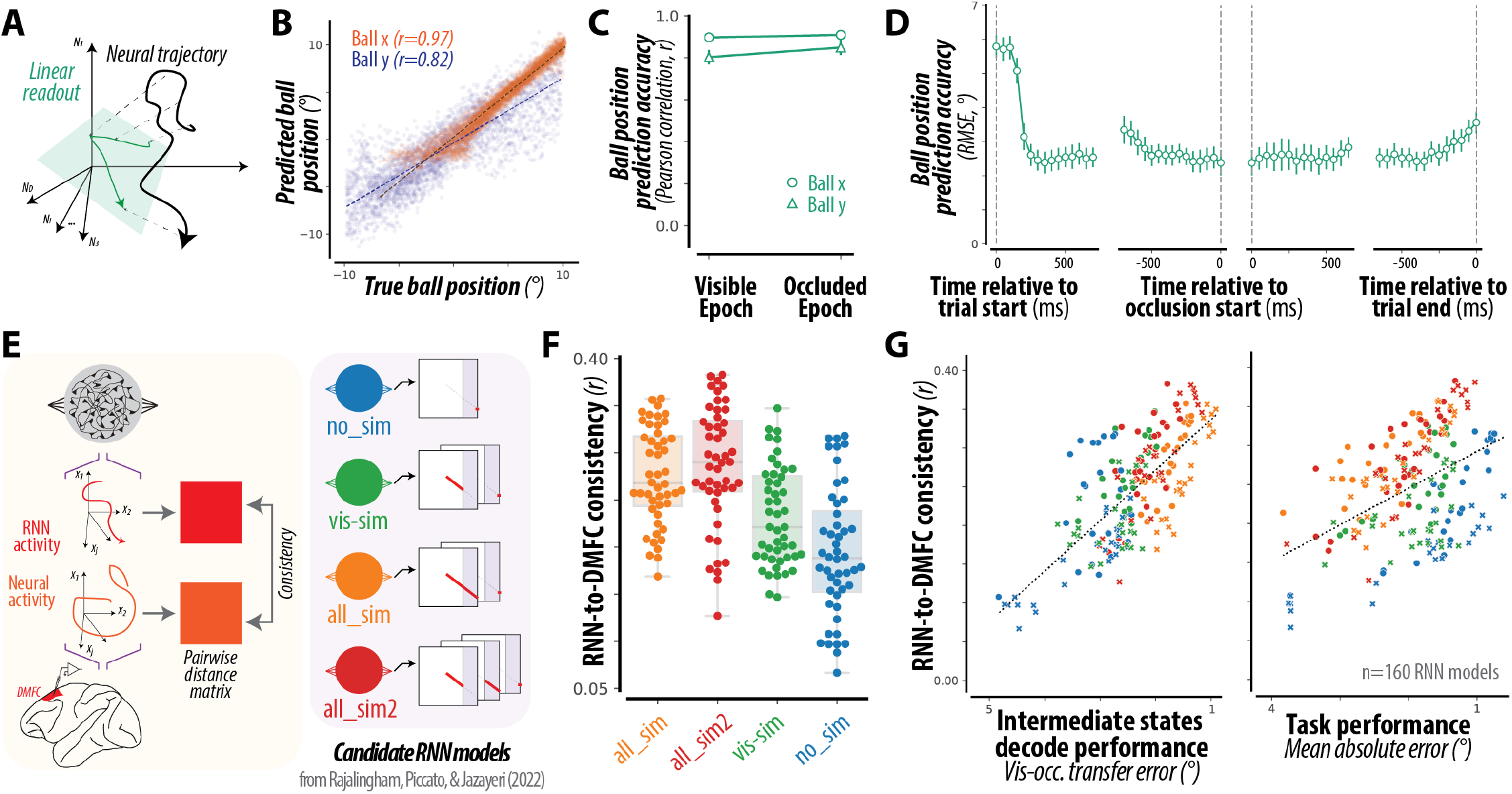
**(A) Conceptual schematic.** To characterize the representation of the latent ball position in the DMFC population, we used cross-validated linear regressions to read out the moment-by-moment position of the ball. The schematic illustrates, for a single Mental-Pong condition, the neural dynamics (black trace), and the corresponding predicted ball position (x, y) (green trace) via a linear readout (green subspace). **(B) Accuracy of DMFC readout**. A linear readout from the DMFC population accurately captures the ball position, as shown by the scatters of true and predicted ball position along horizontal (ball x) and vertical (ball y) dimensions (r=0.97, 0.83, p<10^−100^, p<10^−100^, for ball position x, y respectively). Dotted lines correspond to least squares regression lines. **(C, D) Temporal stability of DMFC readout**. The same linear readout can accurately capture the ball (x,y) position over the entire trial, i.e., (C) over both visible and occluded epochs, as quantified by a Pearson correlation, and (D) over 50ms time-bins throughout the trial, as quantified by an Root-Mean-Squared-Error (RMSE) between true and predicted ball positions. Errorbars denote mean±SE. There was no significant difference in performance between visible and occluded epochs (p>0.05, permutation test of difference in Pearson correlations). **(E) Comparing RNNs to neural dynamics**. (left) To directly compare the internal dynamics of RNNs to the measured neural population dynamics, we first characterized each representation via a metric (pairwise distance matrix), and computed the consistency between corresponding metrics. (right) In prior work, we trained several hundred RNNs, varying in various architectural parameters (e.g., different cell types, number of cells, regularization types, input representation types, etc.), to map visual inputs to an endpoint prediction output, with some RNNs additionally optimized to dynamically track the position of the ball throughout specific trial epochs. The legend illustrates the four different RNN optimization types (no_sim, vis_sim, all_sim, all_sim2), with schematics of their optimization targets. Critically, RNNs were not optimized to reproduce primate behavior, only to solve the task. **(F)** Distribution of DMFC-consistency scores for all RNNs, grouped by optimization types; the swarm plot shows individual models, and the boxplot shows the median, 1st and 3rd quartiles, and range of each distribution. RNNs optimized for dynamic inference better matched DMFC dynamics than RNNs optimized for task performance alone (*p*<*10*^*-8*^ for all_sim vs no_sim, *p*<*10*^*-9*^ for all_sim2 vs no_sim, two sample t-test). **(G) Functional correlates of RNN-to-DMFC consistency**. Across all RNNs, scatter of consistency with DMFC dynamics versus task intermediate state decode performance (left) and task performance (right). The variation in consistency across different RNNs weakly depends on overall task performance (*R*^*2*^*=0*.*24, p*<*10*^*-12*^). Instead, consistency was strongly correlated with ISDP (*R*^*2*^*=0*.*52, p*<*10*^*-32*^). Note that the abscissas are flipped such that left-to-right corresponds to increasing performance (i.e., decreasing error) and increasing dynamic inference ability (i.e., decreasing ISDP error).

Monkeys typically made hand and eye movements that co-varied with the ball’s position (Figure S2B). While such movements are sometimes presented as evidence of an internal tracking strategy ^19,20^, this brings up the possibility that the observed neural dynamics reflect signals related to the preparation and execution of movements. Indeed, we found that DMFC had a representation of many confounding variables, including hand and eye kinematics, and the time within the trial (Figure S2C). However, accounting for these variables using partial correlations did not dramatically reduce the accuracy of the decoded ball position (Figure S2D). Together, these results suggest that DMFC has a low-dimensional representation of the latent instantaneous ball position, consistent with the explicit inference, that is distinct from other behavioral and latent factors.

Previously, we constructed RNN models to solve the Mental-Pong task either with or without the capacity to explicitly track the latent ball position, and observed that primate behavior was best captured by RNNs endowed with the explicit inference strategy. Since these models serve as direct instantiations of hypotheses related to explicit inference versus implicit mapping, we next sought to directly compare representations in DMFC to representations in these different classes of RNNs. As a first step, we computed pairwise distances between all visited states in both DMFC and RNNs over all Mental-Pong conditions. We then quantified the similarity between DMFC and RNNs by measuring the noise-adjusted correlation between those pairwise distances, which we termed consistency (see Methods, Figure 2E). Note that all RNNs were optimized for task performance and pre-registered ^10^, and were not fit to the neural data. Therefore, a measure of similarity between DMFC and different types of RNNs serve as a strong test of explicit inference versus implicit mapping hypotheses.

RNNs exhibited a broad range of consistency scores, with models optimized for explicit inference being most consistent with signals in DMFC (Figure 2F, S2E). To compare all RNNs on the same footing, we developed a common metric to measure the degree an RNN carries explicit information about the ball position while behind the occluder. The metric, which we refer to as the intermediate state decode performance (ISDP), was computed as the mean absolute error between the true time-varying ball position and the predicted position from a cross-validated linear regression (see Methods). Results revealed a strong relationship between ISDP and consistency with DMFC scores across the RNNs (Figure 2G). Critically, this similarity was not explained by RNN performance (Figure S2F) suggesting that the key factor explaining DMFC dynamics was the presence of an explicit representation of the ball’s intermediate positions. These results were consistent across both monkeys (Figure S4). Taken together, these results suggest that the brain explicitly computes the latent position of the occluded ball, consistent with mental simulation.

Our previous work comparing RNNs to behavior and our current results comparing DMFC dynamics to RNNs are both consistent with the idea that the brain solves Mental-Pong via an online simulation of the instantaneous position of the ball. However, we wondered if animals additionally compute a rapid estimate of the ball trajectory shortly after the ball begins to move. Such an early estimate would be advantageous computationally as it could enable animals to plan in advance and initiate paddle movement early in the trial. To test this possibility, we applied a static cross-validated linear regression to neural activity immediately after trial onset (see Methods, Figure 3A). Strikingly, we found an accurate and sustained representation of the interception point that emerged shortly after ball movement onset, long before the ball reached its final position (Figure 3B). This representation was not a trivial consequence of the linear trajectory of the ball since an identical cross-validated linear regression applied to DMFC’s estimates of the instantaneous ball position was unable to predict the interception point with the same level of accuracy (Fig S2G). Moreover, this early estimate was predictive of the animal errors as evidenced by the noise-adjusted correlation across conditions between behavioral errors and errors in the interception point inferred from early DMFC data (Figure 3C, S5). Taken together, these results highlight the intriguing possibility that the online simulation is preceded by computing an estimate of the interception point shortly after the movement onset.

**Figure 3.**
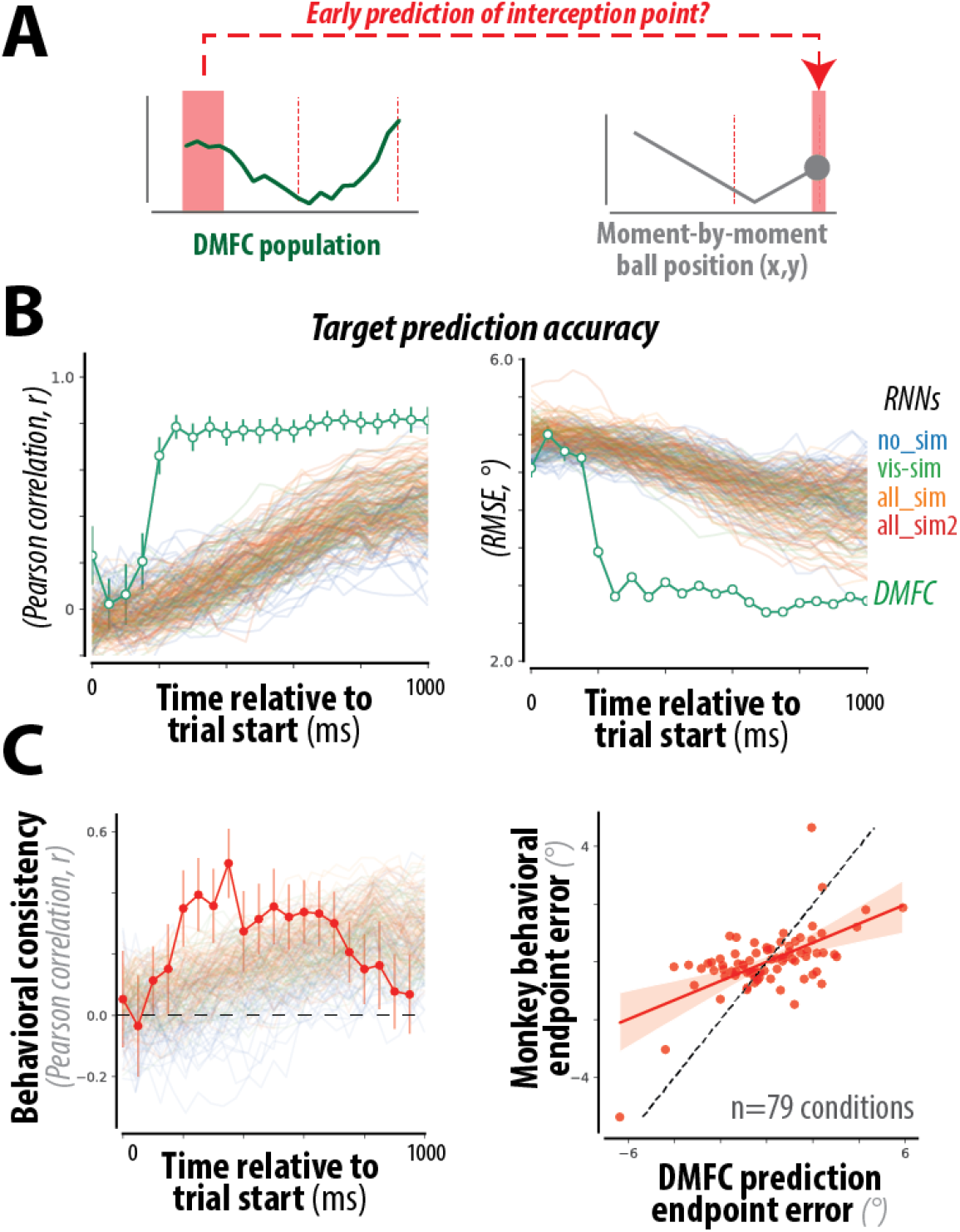
**(A) Conceptual schematic.** We used a cross-validated linear regression to read out the final interception position of the ball from the entire time-course of DMFC activity, and asked whether DMFC exhibits an early prediction of the interception point. **(B) DMFC’s representation of the final ball interception point**. Accuracy of a static read-out of the endpoint ball vertical position over time, quantified via Pearson correlation (left panel) and Root-Mean-Square-Error (RMSE, right panel), for the DMFC population (green) and each of the tested RNNs (colored as in Figure 2E). Errorbars denote mean±SE. To infer statistical significance, performance was compared to a shuffled control (not shown); DMFC predictions were significantly more accurate than the shuffled control for all timepoints following t=250ms (*p*<*10*^*-10*^, permutation test). **(C) Behavioral consistency of DMFC’s interception point estimate**. The behavioral consistency is the noise-adjusted correlation across conditions between DMFC’s interception point representation and the monkeys’ endpoint paddle position, after regressing out the true endpoint ball position from each. (left) DMFC’s early interception point prediction was correlated with the monkeys’ behavioral errors (red, mean±SE). To infer statistical significance, behavioral consistency was compared to zero; DMFC predictions were significantly consistent for all timepoints following t=250ms (*p*<*10*^*-10*^, permutation test). The colored lines show the mean behavioral consistency for each of the tested RNN models (colored as in Figure 2E). (right) Comparison of the endpoint error (residual after regressing out the true endpoint ball position) of the DMFC interception point estimate vs. the monkeys’ behavior, for all 79 conditions at t=400 ms after trial start.

In contrast to DMFC, none of the RNNs exhibited a rapid and sustained representation of the interception point (Figure 3B) and were not predictive of behavioral errors early in the trial (Figure 3C). This discrepancy motivates future models of mental simulation to be appropriately constrained so that they can combine early prediction with online simulation.

## Discussion

The structure and dynamics of population activity in DMFC during Mental-Pong and their similarity to RNNs trained to track the instantaneous ball position suggest that monkeys may mentally track the occluded ball. These findings take an important step toward validating cognitive theories about the role of mental simulation during inference and planning. They also highlight a link between mental simulation and low-dimensional task-relevant neural dynamics in areas of the frontal cortex implicated by prior neuroimaging studies in humans ^15 21–23^.

We here focus on a defining “dynamic inference” feature of mental simulation – that the brain reproduces the neural states associated with *deterministic* physical states – and our results do not speak to *stochastic* simulation approaches that additionally represent state uncertainty (e.g., particle filters) ^2,24–26^. Notably, dynamic inference is not a necessary feature of neural systems that perform Mental-Pong; the target can be computed by a nonlinear function of the ball’s visible state without explicitly tracking. We tested these hypotheses using RNN models optimized to perform Mental-Pong – with or without additional dynamic inference constraints – and, critically, these RNN models were constructed before the collection of any of the neural data, and are pre-registered models of neural dynamics.

Interestingly, DMFC dynamics were consistent with both dynamic inference and direct mapping strategies, suggesting that the brain may use computations at multiple time-scales – both rapid “offline” predictions and dynamic “online” tracking – to support physical inferences via interceptive movements ^27,28^. This heterogeneity of solutions could explain the divergent observations regarding the need for “simulation” in making physical inferences, and as such could help reconcile debates regarding the algorithms underlying our physical inference abilities. We speculate that the presence of both online and offline strategies might reflect a control machinery that computes initial state estimates through a rapid feedforward computation that can be refined using online recurrent computations. Future research is needed to test how such an integrative process is instantiated via neural population dynamics in the fronto-parietal circuit and how different roles each node of the circuit plays. Our work opens a door for a deeper understanding of the neural circuit dynamics underlying physical reasoning.

## Methods

### Subjects and surgery

Two adult monkeys (Macaca mulatta), one male (M) and one female (P), participated in the experiments. For each animal, a surgery under general anesthesia and using sterile surgical technique was performed to implant three pins for head restraint. A second surgery under general anesthesia and using sterile surgical technique was performed to implant a customized rectangular recording chamber over a craniotomy targeting the dorsomedial frontal cortex in the left hemisphere from the top of the skull (Monkey P:

+24 posterior-anterior, 0 medial-lateral; Monkey M: +24 posterior-anterior, 0 medial-lateral). The Committee of Animal Care at Massachusetts Institute of Technology approved the experiments. All procedures conformed to the guidelines of the National Institutes of Health.

### Behavioral task

In Mental-Pong, the player controls the vertical position of a paddle along the right edge of the screen to intercept a ball as it moves rightward. On each trial, the ball starts at a random initial position (x_0_,y_0_) and a random initial velocity (dx_0_, dy_0_), and moves at a constant speed throughout the trial. The screen additionally contains a large rectangular occluder right before the interception point such that the ball’s trajectory is visible only during the first portion of the trial. Trial conditions were constrained by the following criteria: 1) the ball always moved rightward (dx>0), 2) the duration of the visible epoch was within a fixed range ([15,45] RNN timesteps or [624.9, 1874.7] ms), 3) the duration of the occluded epoch was within a fixed range ([15,45] RNN timesteps or [624.9, 1874.7] ms), 4) the number of times the ball bounced was within a fixed range ([0,1]). We previously sampled 200 conditions and reported their stimulus parameter distributions for this task in a large-scale behavioral comparison of human, monkey, and RNN across 200 unique conditions ^10^. Here, we subsampled 79 conditions (26 conditions with a bounce) from this same set of conditions for neurophysiology experiments in monkeys. Stimuli and behavioral contingencies were controlled by an open-source software (MWorks; http://mworks-project.org/) running on an Apple Macintosh platform.

Each trial initiated when the animals acquired and held gaze on a central fixation point (white circle, diameter: 0.5 degree in visual angle in size) within a window of 4 degree of visual angle for 200 ms. Following this fixation acquisition, animals were allowed to make any eye movements and freely view the screen. Afterwards, the Mental-Pong condition was rendered onto the screen with the entire frame spanning 20 degrees of visual angle: the ball was rendered at its initial position (x_0_, y_0_), and the paddle was rendered in the central vertical position at the right edge of the frame. 300ms after this framer render, the trial began with the ball moving in its initial velocity (dx_0_, dy_0_); this pre-trial delay was included to mitigate potential visually-driven transients in the neural recordings.

As shown in Figure 1A, the paddle was initially rendered as a small, transparent green square (0.5 deg × 0.5 deg), but turned into a full paddle (0.5 deg × 2.5deg) when the animals first initiated paddle movements. This feature enforced that animals performed a movement (i.e., trigger the paddle) on all trials. For the remainder of the trial, animals could freely move the paddle up or down using a customized joystick. The paddle position was updated on every screen refresh (i.e., every 16.6ms), and moved at a constant speed of 0.17deg/16ms = 0.01deg/ms. Trial ended when the ball reached the right-end of the screen. Upon the trial’s end, the occluder disappeared to give animals feedback of their performance. If they successfully intercepted the ball, it would bounce off their paddle (see Figure 1A); if they had failed to intercept it, it would continue its path off the frame. Trials were separated by an inter-trial-interval of 750 ms.

In prior work, we found that monkeys rapidly learn to perform this task and immediately generalize to novel task conditions, suggesting that Mental-Pong taps into pre-existing behavioral abilities of macaque monkeys and, as such, is particularly suitable for neurophysiology experiments in monkeys.

### Monkey behavior

During the experiments, monkeys were seated comfortably in a primate chair, and were head-restrained. For training purposes, we first acclimated monkeys to a 1 degree-of-freedom joystick placed right in front of the primate chair. Next, we started a curriculum for training animals to perform Mental-Pong. Monkeys were first trained to use the joystick to control the vertical position of a paddle, and then practiced Mental-Pong using 200 unique trial conditions. We previously reported that monkeys rapidly learn to perform Mental-Pong and generalize to novel conditions on the very first trial, suggesting that this task taps into pre-existing physical inference abilities^10^. Moreover, monkeys have remarkably human-like patterns of endpoint paddle error over conditions, suggesting that they are a suitable model of human physical inference abilities.

For all experiments, the stimuli were presented on a fronto-parallel 23-inch (58.4-cm) monitor at a refresh rate of 60 Hz. Eye position was tracked every 1 ms with an infrared camera (Eyelink 1000; SR Research Ltd, Ontario, Canada). The joystick voltage output (0-5V) was converted to one of three states (up: 3-5V; down: 0-2V; neutral: 2-3V), which was used to update the position of the paddle.

### DMFC Neurophysiology

We recorded the activity of neural populations from the dorsomedial frontal cortex (DMFC) in two monkeys while they performed Mental-Pong. To visualize and plan the anatomical location of recording sites, we first performed a structural MRI scan where a recording grid filled with a mixture of agarose and gadolinium was inserted into the chamber. We recorded neural activity in the dorsomedial frontal cortex (DMFC) with 64-channel linear probes (Plexon V-probes, monkey P) and high-density silicon probes (Neuropixels, monkey M). Recording locations targeted the region of cortex located anterior to the genu of the arcuate sulcus, and medial to the upper arm of the arcuate sulcus. We conservatively refer to this region as dorsomedial frontal cortex (DMFC), potentially comprising distinct functional regions, e.g. supplementary eye field, dorsal supplementary motor area (SMA), and pre-SMA (no recordings were made in the medial bank). In monkey P, we systematically sampled DMFC by recording from all grid locations spaced 1 mm apart within an 8 mm × 3 mm region (24 distinct recording sites, each sampled in two sessions). In monkey M, we recorded from a subset of grid locations spaced 1 mm apart within a 8 mm × 1 mm region (6 distinct recording sites, each sampled in one session). To bolster the reproducibility of the neural recording process, we did not select recording locations based on the existence of task-relevant modulation of neural activity, but instead systematically sampled the cortical tissue. Recording sites are shown in Figure S1.

### Neural data pre-processing

We used KiloSort (KS) 3.0 to sort the spike waveform data, using all default parameters with the exception of *ops*.*Th=[9,4]*. We did not perform any post-hoc manual resorting of spike waveforms. Instead, we used a custom Python library for post-processing based on the Kilosort-outputed spike templates, which performed removal of unstable units and merging of similar KS templates into units. This processing step acted largely as a filter: ∼80% of output units were sorted as per KS 3.0 alone. We then visually inspected all recording sessions using summary figures for waveform similarity. Our goal was to ensure reproducibility of the spike sorting process by using a fully-automated sorting process, while reserving human intervention strictly for inspection and validation.

For each recording session, we averaged the spike trains (at 1 ms resolution) across repetitions of the same task conditions, including both success and failure trials. This resulted in a data matrix of size *N*_*units*_ *× N*_*condition*_ *× N*_*time*_ for each session, where *N*_*condition*_*=79, and N*_*time*_*=5000 ms*. Given the large number of unique conditions (a total of 79 × 2=158 conditions, over “visible” and “occluded” task settings), and given the stability of recorded units, not all neurons were recorded during all possible conditions. We excluded neurons in which more than 5 Mental-Pong conditions were missing. For the remaining neurons, we imputed any missing values using standard matrix imputation, replacing missing entries with the global mean of the neuron’s firing rate (across time and conditions). At this threshold, less than 1% of data bins were imputed.

We then pooled all sessions together, and downsampled the trial-averaged spike responses by binning into independent time-bins of 50ms. To avoid introducing spurious temporal dependencies, we performed no temporal smoothing of the binned spike data. The total number of recorded neurons was 2058, 518 for monkeys P and M, respectively.

We next sought to exclude units with unreliable response patterns (e.g. due to noise, low number of trials, etc.). For each unit, we randomly split all trials into two equal halves and estimated the response matrix from each half, resulting in two independent estimates of the unit’s response pattern of size *N*_*condition*_ *× N*_*time*_. We took the Pearson correlation between these two estimates as a measure of the reliability of that neural response given the amount of data collected, i.e., the split-half internal reliability. We then excluded all units where the split-half reliability was not statistically significant (p>0.05). Note that this selection process does not select for neurons modulated by task variables, but only for neurons with reliable response patterns. Finally, this resulted in a data matrix of size of size *N*_*units_all*_ *× N*_*condition*_ *× N*_*timebins*_. The total number of reliably recorded neurons was *N*_*units_all*_ = 1889 (1552, 337 for monkeys P and M, respectively).

We found that neural responses of *N*_*units_all*_ could be captured by a smaller number of dimensions (Figure 2). Thus, we used Factor analysis (with *n=50* factors) to capture variability in neural activity that is shared across neural populations ^29^, resulting in a population response matrix of *N*_*factors*_ *× N*_*condition*_ *× N*_*timebins*_. Given that the factors are not necessarily orthonormal, we additionally performed both with and without VARIMAX alignment, with no difference in decoding results.

### Decoding analyses

To test the capacity of neural populations to perform dynamic tracking, we used cross-validated linear regressions to read out specific task/behavioral variables (e.g. the × and y position of the ball over time) from the neural population data. Regressions were performed using the scikit-learn python library. For all analyses, we focused on “static” read-outs, whereby a single regression was learned to map all time points of neural activity to the time-varying task variable. To do this, we reshaped the neural population data by pooling along condition and time dimensions, resulting in a matrix of size (*N*_*cond*_*N*_*timebins*_) × *N*_*factors*_. However, we did not consider each row of this data as an independent observation. Instead, regressions were trained and tested on distinct Mental-Pong conditions (using GroupShuffleSplit). We measured the goodness of regressions using several metrics: Pearson correlation between the true and predicted variable, and the root mean squared error. For all analyses (except Figure S2A), cross-validated regressions were trained on 50% of the conditions, and repeated over 100 train/test splits.

To account for possible covariations due to sensorimotor variables, we also computed partial Pearson correlations, the estimated correlation after regressing out co-varying attributes (candidate sensorimotor variable, e.g. eye position x) with a linear least squares regression.

In one analysis, we use DMFC’s estimate of the ball position as the features for regressing the final ball interception point. To do so, we first trained and tested linear regressions to predict the ball position from DMFC activity; this procedure was repeated over 100 train/test splits, and ball position predictions were averaged across these. The average ball position predictions were then used to train and test linear regressions predicting the final ball interception point.

### RNN models

To yield insight into the computations that generate these neural dynamics, we directly compared DMFC dynamics with the corresponding dynamics of recurrent neural network (RNN) models. In previous work^10^, we constructed several hundreds of RNN models optimized to perform the same task as humans and monkeys. RNNs were trained to map a series of visual inputs (pixel frames) to a movement output, where the target movement output corresponded to a prediction of the particular paddle position at a particular time point in order to intercept the ball. Different RNN models varied with respect to architectural parameters (different cell types, number of cells, regularization types, input representation types), and were differently optimized (one of four different target outputs, either with or without dynamic inference). Critically, RNNs were not optimized to reproduce primate behavior, only to solve specific tasks.

Critically, we trained RNNs with one of four different optimization types, which we code-named as no_sim, vis-sim, all_sim and all_sim2. For all RNN types, one of the output channels (called “movement output”) corresponded to the paddle position, which was optimized to predict only two samples per trial: one consisting of the initial central paddle position, and the second corresponding to the particular paddle position at a particular time point in order to intercept the ball. As shown in Figure 2E, this was the only loss term for RNNs of the “no_sim” class. For the remaining RNNs, we additionally estimated a loss term from some of the other channels, as the mean squared error between the channel output and a target time-varying signal corresponding to the ball’s position (x,y) during specific trial epochs (“vis-sim”: visual epoch only; “all_sim”: entire trial; “all_sim2”: separate channels for visual and occluded epochs). This set of optimization choices explored different computational constraints regarding the specifics of dynamic inference processes. For instance, “vis_sim” corresponds to constraints on the sensory but not the latent computations. On the other hand, “all_sim” and “all_sim2” correspond to shared and independent constraints, respectively, on the sensory and latent computations.

RNNs were trained using the TensorFlow 1.14 library using standard back-propagation and adaptive hyperparameter optimization techniques ^30^; training each RNN took approximately one day on a Tesla K20 GPU. Importantly, these RNN models are pre-registered models of neural dynamics: they were constructed and tested on behavior alone before the collection of any of the neural data.

### RNN to DMFC comparisons

Consider each representation, whether neural or artificial, as a response matrix of size *N*_*units*_ *× (N*_*cond*_*N*_*timebins*_), where *N*_*cond*_*N*_*timebins*_ is the total number of neural states *N*_*states*_. Each representation can be characterized via a matrix of pairwise distances between all states, resulting in a matrix of size *N*_*states*_ *× N*_*states*_ (Figure 2E). We estimated the similarity (termed *consistency*) between two representations via a noise-adjusted correlation between their corresponding pairwise distances ^31^.

To do so, we first randomly split all trials into two equal halves and estimated the pairwise distance matrix from each half, resulting in two independent estimates of the system’s response. We took the Pearson correlation between these two estimates as a measure of the reliability of that response given the amount of data collected, i.e., the split-half internal reliability. To estimate the noise-adjusted correlation, we computed the Pearson correlation over all the independent estimates of the pairwise distance matrix from the model (**m**) and the brain (**b**), and we then divide that raw Pearson correlation by the geometric mean of the split-half internal reliability of the same pairwise distance matrix measured for each system:

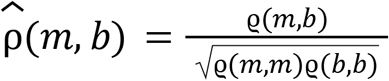

Since all correlations in the numerator and denominator were computed using the same amount of trial data (exactly half of the trial data), we did not need to make use of any prediction formulas (e.g. extrapolation to larger number of trials using Spearman-Brown prediction formula). This procedure was repeated 10 times with different random split-halves of trials. Our rationale for using a reliability-adjusted correlation measure for *consistency* was to account for variance in the pairwise distance matrices that arises from “noise,” i.e., variability that is not replicable by the experimental condition, and thus that no model can be expected to predict. In sum, if the model (**m**) is a perfect replica of the brain (**b**), then its expected consistency score is 1.0, regardless of the finite amount of data that is collected.

This procedure was restricted to measuring distances between states during the occluded epoch, where our previous work showed that RNNs most differ. However, results are similar when considering the full trial (Figure S2E, F).

## Acknowledgments

R.R. is supported by the Helen Hay Whitney Foundation.

H.S. is supported by a NARSAD young investigator grant from the Brain & Behavior Research Foundation.

M.J. is supported by NIH (NIMH-MH122025), the Simons Foundation, the McKnight Foundation, and the McGovern Institute.

## Notes

### Competing Interest Statement

The authors have declared no competing interest.

