## Supplemental Information for "Dynamic tracking of objects in the macaque dorsomedial frontal cortex"

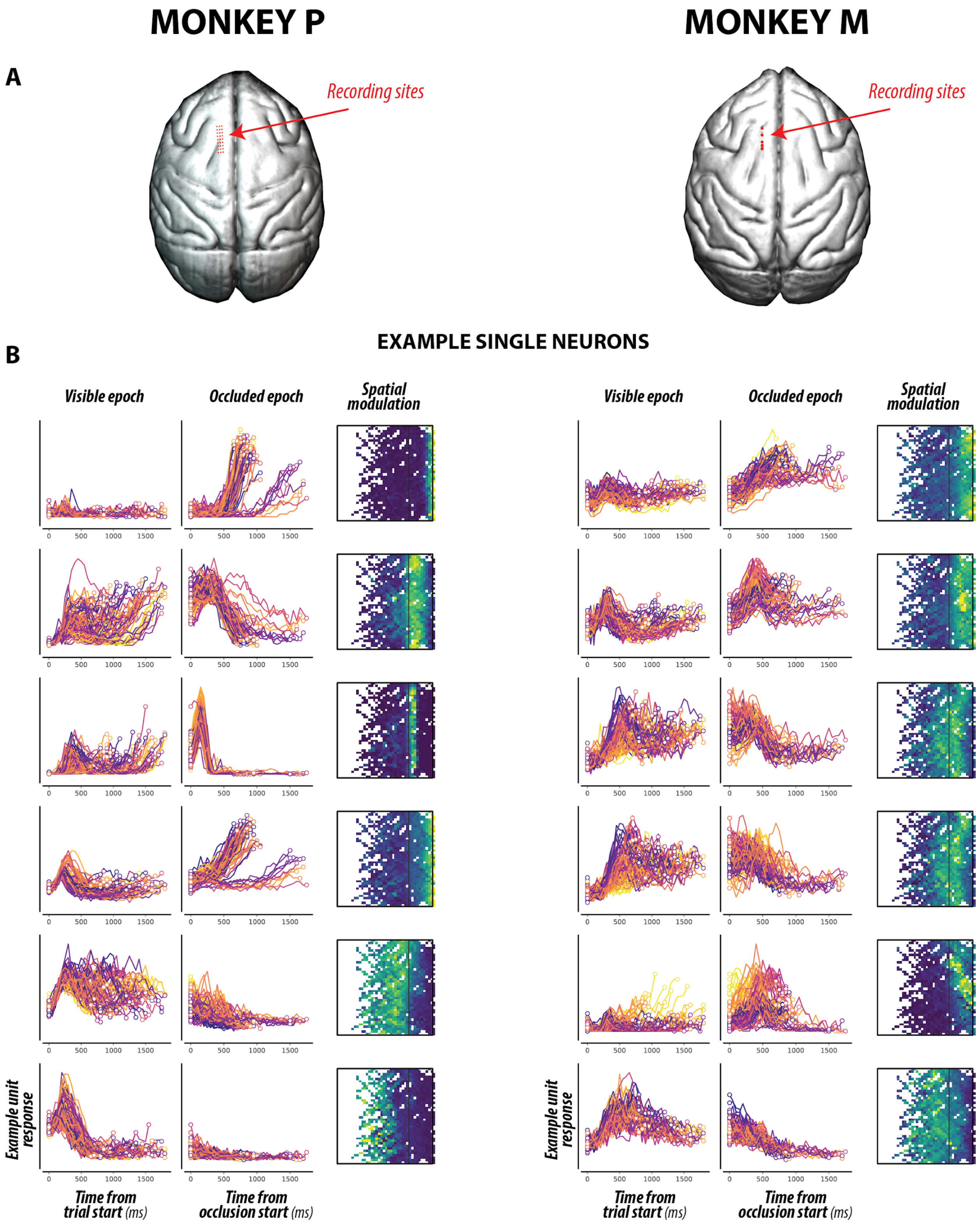


**Figure S1. Neurophysiology. (A)** Neurophysiology sites in DMFC (red circles) rendered on the cortical surface (extracted from structural MRI) of each monkey. **(B)** Activity of six example neurons from each animal. Average responses for each of the 79 M-Pong conditions during the visible epoch aligned to the start of the trial (left), and during the occluded epoch aligned to the start of the occlusion (middle). Color mapping identical to Figure 1C. The right panels show the same activity over the visuospatial dimensions, with the boundary of the M-Pong frame denoted in black.

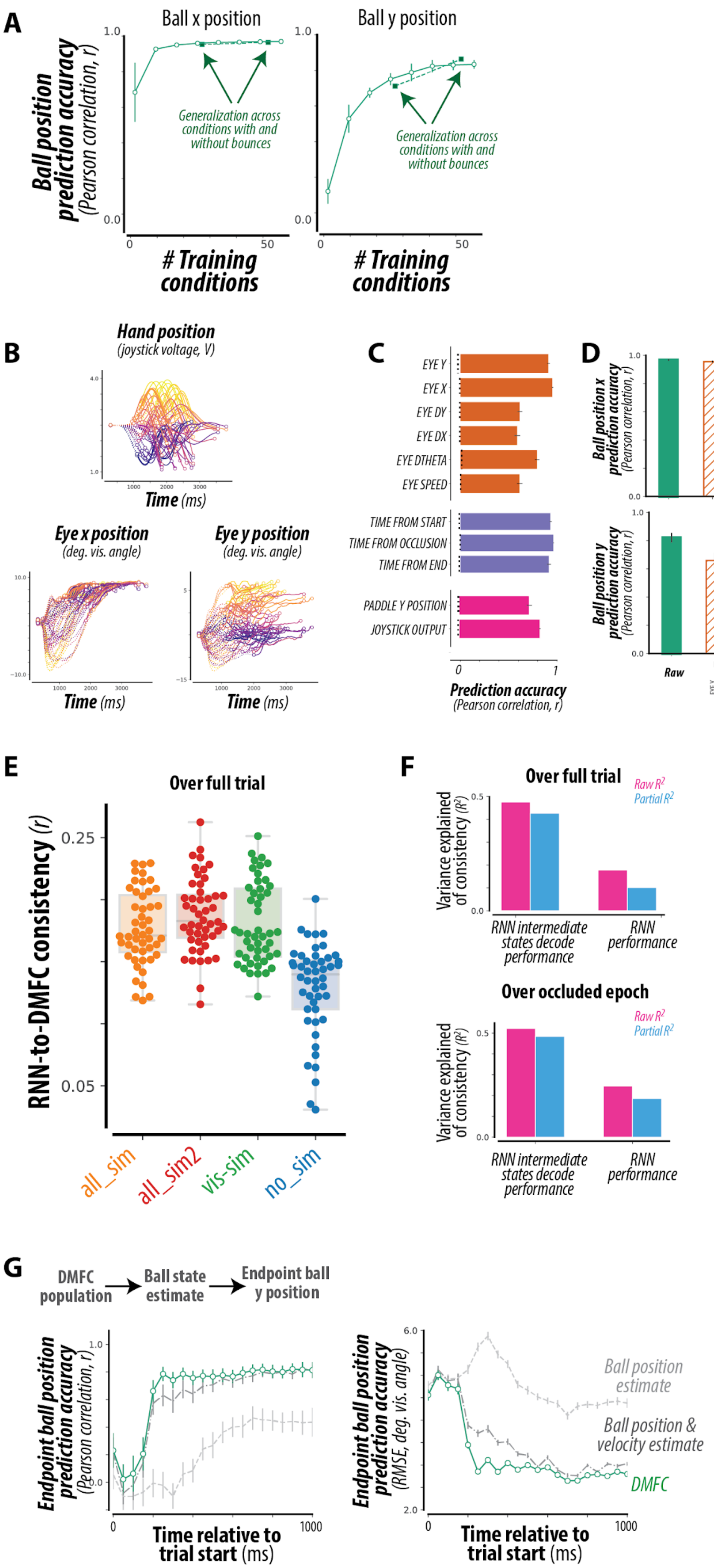
A
Ball x position
Ball y position
Ball position prediction accuracy (Pearson correlation, r)
1.0
0.0
0
50
Generalization across conditions with and without bounces
### Training conditions
B
Hand position (joystick voltage, V)
Time (ms)
Eye x position (deg. vis. angle)
Eye y position (deg. vis. angle)
C
EYE Y
EYE X
EYE DY
EYE DX
EYE DTHETA
EYE SPEED
TIME FROM START
TIME FROM OCCLUSION
TIME FROM END
PADDLE Y POSITION
JOYSTICK OUTPUT
0
1
Prediction accuracy (Pearson correlation, r)
D
Ball position x prediction accuracy (Pearson correlation, r)
Ball position y prediction accuracy (Pearson correlation, r)
Raw
Partial
E
Over full trial
RNN-to-DMFC consistency (r)
0.25
0.05
all_sim
all_sim2
vis-sim
no_sim
F
Over full trial
Over occluded epoch
Raw R2
Partial R2
Variance explained of consistency (R2)
0.5
0.0
RNN intermediate states decode performance
RNN performance
G
DMFC population
Ball state estimate
Endpoint ball y position
Endpoint ball position prediction accuracy (Pearson correlation, r)
Endpoint ball position prediction accuracy (RMSE, deg. vis. angle)
6.0
2.0
1000
Time relative to trial start (ms)
Ball position estimate
Ball position & velocity estimate
DMFC

**Figure S2. Control analyses. (A) Robustness of latent object readout.** Accuracy of the readout of ball position x and y, as a function of the number of Mental-Pong conditions used for training. Read-outs could accurately generalize across conditions from training on a smaller number of conditions. The green squares indicate generalization from training on conditions with bounces and testing on conditions without bounces, and vice versa. Errorbars denote mean±SE. **(B) Hand and eye movements.** Average time-courses of the hand and eye position across all M-Pong conditions, with color mapping identical to Figure 1C. **(C) Representation of sensorimotor variables in DMFC population.** Using the same cross-validated linear read-out approach as in Figure 2B, DMFC accurately represented many sensorimotor variables, including the movement kinematics (position, velocity, and acceleration) of the hand and eye, and the time within trial relative to the start, middle, and end of the trial. Partial correlations are significantly greater than chance for all tested covariate ($r>0.6$, $p<10^{-100}$ for all bars). **(D) Accounting for sensorimotor variables.** Given the reliable differences between sensorimotor variables and ball position**,** the accuracy of the latent object readout is shown after accounting for each sensorimotor variable, via a partial correlation. **(E)** Distribution of human-consistency scores for all RNNs, grouped by optimization types as in Figure 2F, but when comparing RNNs and DMFC over the entire trial. The swarm plot shows individual models, and the boxplot shows the median, 1st and 3rd quartiles, and range of each distribution. **(F) Functional correlates of RNN-to-DMFC consistency.** Quantifying the relationship shown in Figure 2G for both the entire trial (top), and for the occluded epoch only (bottom). The strength of dependence between functional attributes and consistency with DMFC dynamics is shown as a proportion of variance explained ($R^2$, pink bars). Partial $R^2$ (blue bars) measures this strength after accounting for covariations due to the other attribute. Across all RNNs, consistency with DMFC dynamics was well explained by intermediate state decode performance ($R^2=0.47$, $0.52$, $p<10^{-28}$, $p<10^{-32}$ for top and bottom panels respectively) and also weakly explained by overall task performance ($R^2=0.18$, $24$, $p<10^{-9}$, $10^{-13}$ for top and bottom panels respectively). **(G) Comparison of DMFC representation of target with control hypothesis.** Accuracy of different read-outs of the endpoint ball y-position (the target), quantified via Pearson correlation (left) and error (right). We first trained a read-out of the DMFC population to predict the moment-by-moment ball state. From this ball state estimate, we trained a read-out to predict the target. The light gray line shows the accuracy of this target estimate, when the ball state corresponds to the ball position (x,y). The dark gray line shows the corresponding accuracy when the ball state corresponds to the ball position and velocity (x, y, dx, dy). The accuracy of the read-out of the target directly from the DMFC population (green) is greater than each of these controls ($p<0.001$ for all timepoints in [250ms, 500ms] for RMSE).

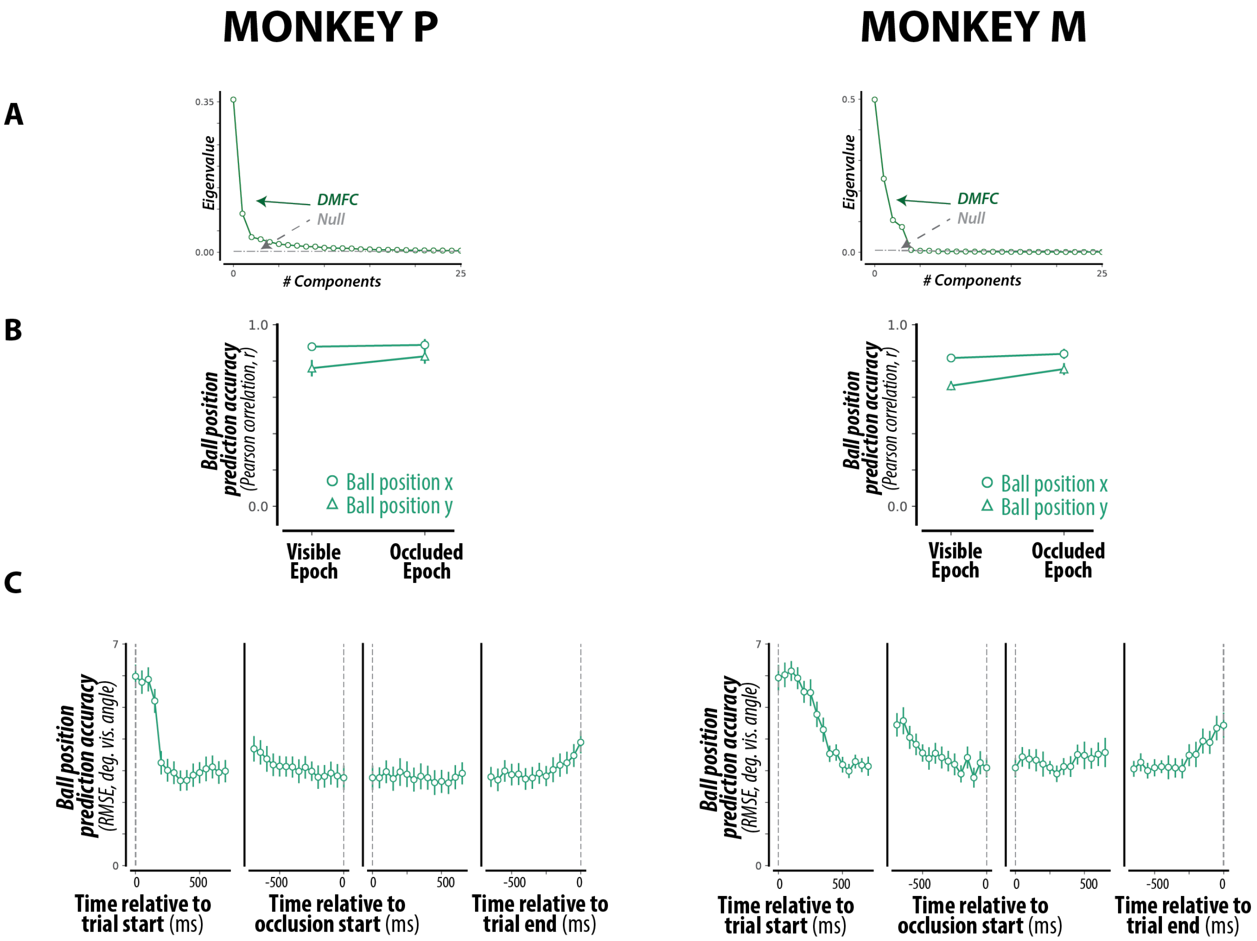


**Figure S3. Latent object readout. (A)** Neural dynamics in DMFC were relatively low-dimensional, as quantified by the eigenspectrum, for both animals. **(B)** Accuracy of latent object read-out over visible and occluded epochs, quantified by a Pearson correlation, for both animals. **(C)** Accuracy of latent object read-out over 50ms time-bins throughout the trial, as quantified by an RMSE, for both animals.

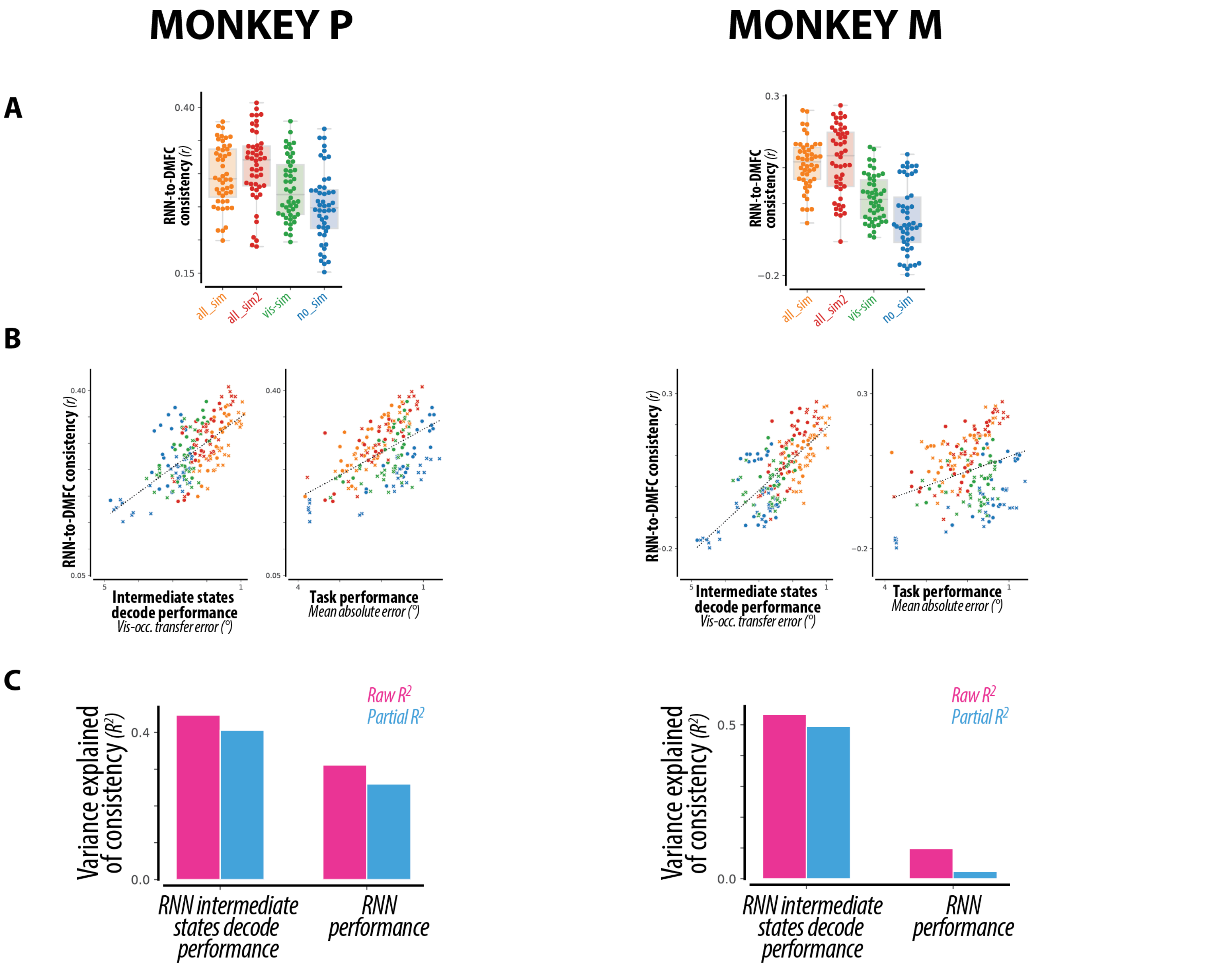


**Figure S4. RNN to DMFC comparisons. (A) RNN-to-DMFC consistency scores.** Distribution of DMFC-consistency scores for all RNNs, grouped by optimization types, for each monkey. The swarm plot shows individual models, and the boxplot shows the median, 1st and 3rd quartiles, and range of each distribution. **(B) Functional correlates of RNN-to-DMFC consistency.** Across all RNNs, scatter of consistency with DMFC dynamics versus task intermediate state decode performance (left) and task performance (right), for each monkey. The variation in consistency across different RNNs weakly depends on overall task performance. Instead, consistency was strongly correlated with ISDP. Note that the abscissas are flipped such that left-to-right corresponds to increasing performance (i.e., decreasing error) and increasing dynamic inference ability (i.e., decreasing ISDP error). **(C) Functional correlates of RNN-to-DMFC consistency.** Quantifying the relationship shown in (B): the strength of dependence between functional attributes and consistency with DMFC dynamics is shown as a proportion of variance explained ($R^2$, pink bars). Partial $R^2$ (blue bars) measures this strength after accounting for covariations due to the other attribute. Across all RNNs, consistency with DMFC dynamics was well explained by intermediate state decode performance, and also weakly explained by overall task performance.

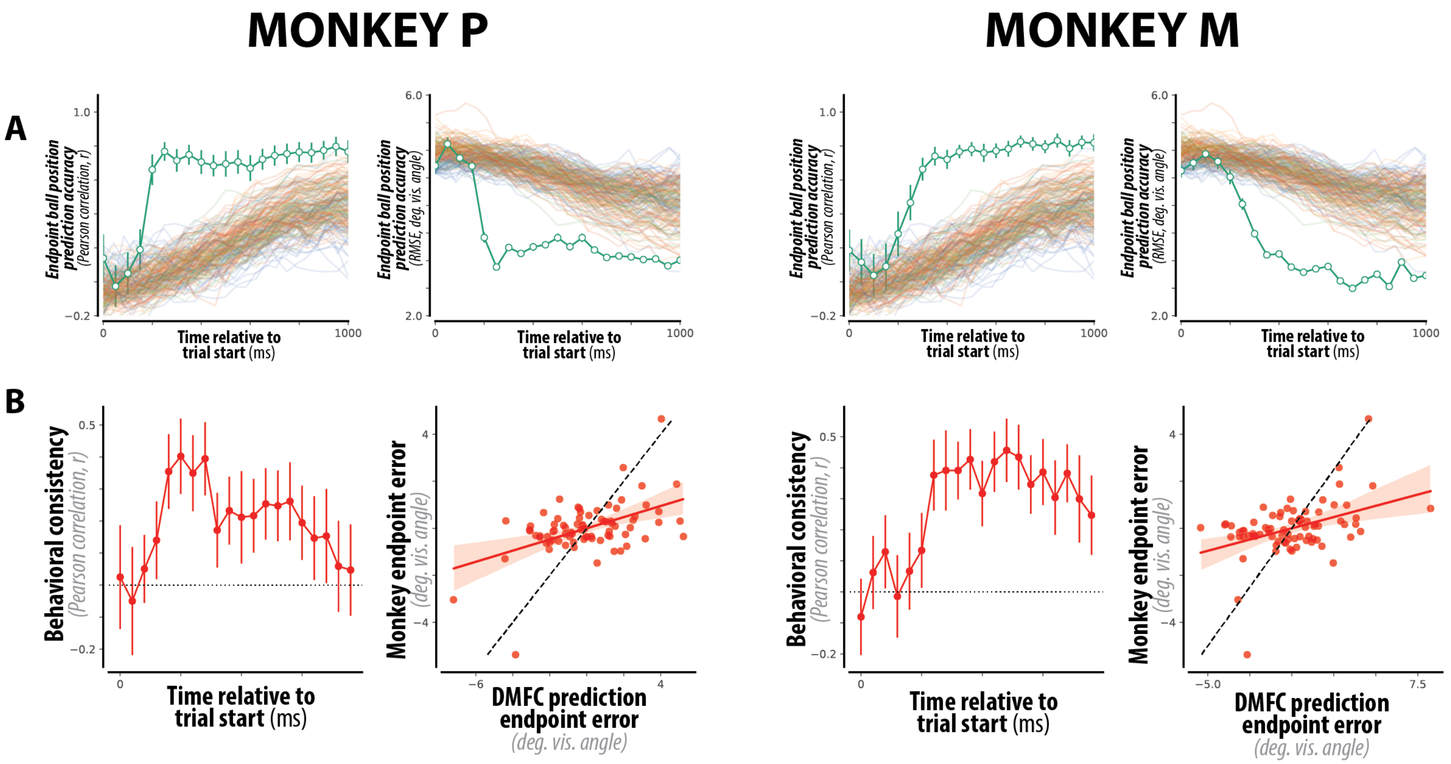


**Figure S5. Rapid target representation. (A) DMFC's target representation.** For each monkey, accuracy of a static read-out of the endpoint ball position over time, quantified via Pearson correlation (left panel) and error (right panel), for the DMFC population (green) and each of the tested RNNs (colored). **(B) Behavioral consistency of DMFC's target representation.** For each monkey, the behavioral consistency is the noise-adjusted correlation across conditions between DMFC's target representation and the monkeys' endpoint paddle position, after regressing out the true endpoint ball position from each. (left) DMFC's early target prediction was correlated with the monkeys' behavioral errors. (right) Comparison of the endpoint error (residual after regressing out the true endpoint ball position) of the DMFC target prediction vs. the monkeys behavior, for all 79 conditions at t=400ms after trial start.
